# Development and validation of an AI/ML platform for the discovery of splice-switching oligonucleotide targets

**DOI:** 10.1101/2022.10.14.512313

**Authors:** Alyssa D Fronk, Miguel A Manzanares, Paulina Zheng, Adam Geier, Kendall Anderson, Vanessa Frederick, Shaleigh Smith, Sakshi Gera, Robin Munch, Mahati Are, Priyanka Dhingra, Gayatri Arun, Martin Akerman

## Abstract

This study demonstrates the value that artificial intelligence/machine learning (AI/ML) provides for the identification of novel and verifiable splice-switching oligonucleotide (SSO) targets *in-silico*. SSOs are antisense compounds that act directly on pre-mRNA to modulate alternative splicing (AS). To leverage the potential of AS research for therapeutic development, we created SpliceLearn™, an AI/ML algorithm for the identification of modulatory SSO binding sites on pre-mRNA. SpliceLearn also predicts the identity of specific splicing factors whose binding to pre-mRNA is blocked by SSOs, adding considerable transparency to AI/ML-driven drug discovery and informing biological insights useful in further validation steps. Here we predicted *NEDD4L* exon 13 (*NEDD4Le13*) as a novel target in triple negative breast cancer (TNBC) and computationally designed an SSO to modulate *NEDD4Le13*. Targeting *NEDD4Le13* with this SSO decreased the proliferative and migratory behavior of TNBC cells via downregulation of the TGFβ pathway. Overall, this study illustrates the ability of AI/ML to extract actionable insights from RNA-seq data. SpliceLearn is part of the SpliceCore® platform, an AI/ML predictive ensemble for AS-based drug target discovery.

## Introduction

RNA splicing is the mechanism by which introns are removed from newly transcribed pre-mRNA to produce mature mRNA. Alternative splicing (AS), a process by which exonic sequences are differentially skipped or included in the mRNA, allows multiple protein isoforms to be encoded by a single gene. AS regulates many biological processes such as cell differentiation, cell state reprogramming, and stress response (Ule & Blencowe, 2019). AS also plays a role in cancer, where tumor-specific AS events can drive tumor progression, metastatic transition, and drug resistance, among other hallmarks of cancer (Urbanski et al., 2018). The growing body of evidence reinforcing the importance of AS in cancer highlights an opportunity to target and drug AS events. While AS can drastically change the structure and function of the resulting protein, it is often difficult to design novel treatments to specifically drug AS protein isoforms. Conversely, modulation of RNA splicing represents a powerful tool for targeting ‘undruggable’ targets, as these treatment modalities can act directly on mRNA or pre-mRNA (Havens & Hastings, 2016).

Splice-switching oligonucleotides (SSOs) are effective in blocking the interaction between splicing factors (SFs) and their pre-mRNA targets in the nucleus to change AS outcomes.(Havens & Hastings, 2016) SSOs can target mis-splicing or disease-specific AS events ahead of translation, allowing the SSO to potentially modulate downstream protein activity without the need to inhibit protein function.(Havens & Hastings, 2016) Recent advances in antisense chemistry and delivery have led to successful clinical translation of oligonucleotide drugs in muscle and the spinal cord diseases (Centa et al., 2020; Finkel et al., 2017; Han et al., 2020; J. Kim et al., 2019; Syed, 2016; K. R. Wagner et al., 2021). While SSOs for monogenic CNS diseases have shown success in clinical trials, most antisense oligonucleotides currently under investigation are for the treatment of cancer (H. Xiong et al., 2021), including but not limited to SSOs targeting AR-V7, PKM, and BCL-X pre-mRNAs (Z. Li et al., 2016; Ma et al., 2021; Yamamoto et al., 2015). To unlock the innovative potential of SSO treatments, there is a need for efficient, scalable technologies to discover novel drug targets and develop SSO compounds for the treatment of cancer and other diseases in which AS mis-regulation plays a key role (Dvinge et al., 2016; Kahles et al., 2018; Park et al., 2019; Urbanski et al., 2018).

In this study, we utilized the SpliceCore® platform (https://www.envisagenics.com/platform/), an ensemble of artificial intelligence/machine learning (AI/ML) algorithms for the discovery of novel therapeutic SSOs based on the analysis of RNA-seq data. We demonstrate the utility of this platform by discovering a novel AS target in Triple Negative Breast Cancer (TNBC), followed by SSO design and validation. We highlight the use of SpliceLearn™, the newest AI/ML algorithm from the SpliceCore platform, to identify AS regulatory elements amenable to SSO targeting. SpliceLearn was trained using sequence-specific binding profiles of SFs along with spliceosome assembly information based on SF-RNA and SF-SF interactions. As a result, SpliceLearn not only predicts the optimal binding position of SSOs that modulate AS, but also informs the identity of SF regulatory networks under steric inhibition on a given RNA, allowing for more transparent and actionable SSO predictions.

The primary goal for this study was to leverage an AI/ML approach to efficiently analyze RNA-seq data for the identification of AS targets amenable to SSO modulation. Once identified, our secondary goal was to develop and experimentally validate a new SSO compound to modulate the AS in TNBC models. We present experimental evidence showing that a novel target discovered with SpliceCore, *NEDD4L* exon 13 (*NEDD4Le13*), was successfully targeted with SSOs predicted with SpliceLearn to promote cell death specifically in TNBC by modulating the TGFβ pathway. Overall, these data lend credence to the use of AI/ML for drug discovery by identifying innovative yet verifiable drug targets *in-silico*, and provide evidence for a novel therapeutic candidate for TNBC.

## Results

### Development of the SpliceCore ensemble of AI/ML algorithms for target discovery

SpliceCore is Envisagenics’ software platform that applies AI/ML algorithms to RNA-seq data to discover disease-specific and biologically relevant AS events amenable to SSO modulation (Figure 1A). To demonstrate the unique potential of SpliceCore in identifying new drug targets, we sought to investigate the cancer genome atlas (TCGA), one of the most popular sources of RNA-seq data in cancer studies (Koboldt et al., 2012). Our premise was to accurately identify a novel SSO drug target in a highly accessed dataset like TCGA, to prove that SpliceCore can discover new targets and extract value from public data, even if it has been used thousands of times in the past.

**Figure 1.**
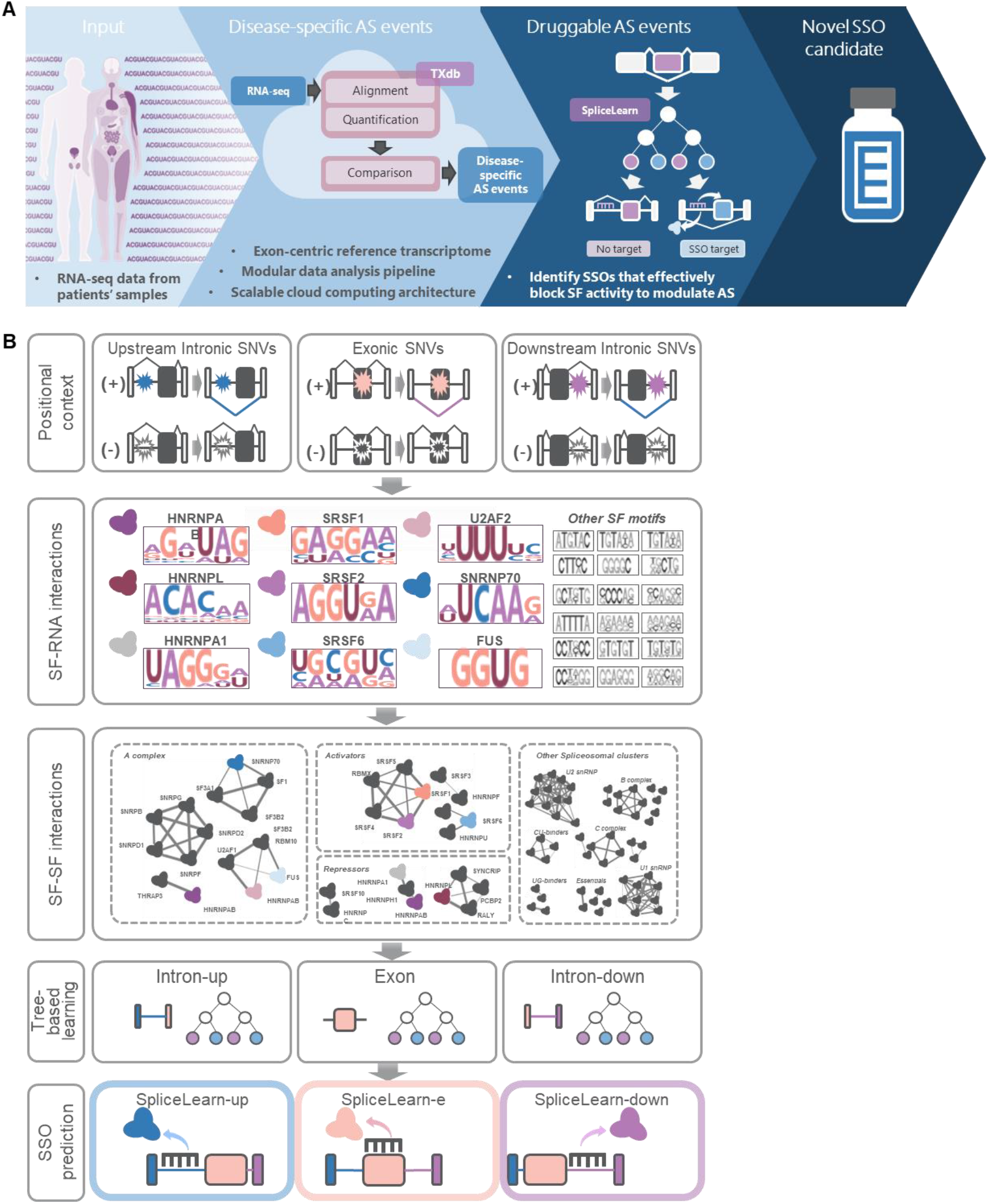
Overview of SpliceCore and SpliceLearn. A. Overview of the SpliceCore platform, including SpliceLearn. Using RNA-seq data as its input, the SpliceCore platform performs de-novo transcript assembly using an exon-centric reference transcriptome called TXdb. It integrates several AI/ML algorithms that allow for modular data analysis and is implemented on the Microsoft Azure cloud to efficiently scale resources. SpliceLearn takes disease-specific AS events identified by SpliceCore and uses a novel AI/ML algorithm to identify functional binding sites for SSOs. B. SpliceLearn was trained on splicing regulatory information including SF binding profiles to RNA as well as SF-SF interactions using tree-based learning for upstream introns, exons and downstream introns independently.

In brief, SpliceCore performs *de-novo* transcript assembly using an exon-centric reference transcriptome called TXdb (Wu et al., 2011). Exon-centric analysis differs from transcript-centric analysis in that it treats the transcriptome as a collection of independent AS events rather than full-length transcripts (Supplemental Figure 1A-B, Methods). Exon-centric analysis is best suited for SSO target discovery because these compounds operate at short sequence length to modulate the inclusion or skipping of a target exon.

TXdb assembly produced a total of 1,743,426 unique AS events derived from 1,252 breast cancer RNA-seq samples in TCGA, where 68% of the assemblies were cassette exon trios, and the remaining 32% were divided among other AS types (Supplemental Figure 1C). In addition, only 4% of the assemblies belonged to previously known AS events, 31% were “supported” assemblies (i.e., constitutive exons) and 65% were novel assemblies (Supplemental Figure 1D).

Following TXdb assembly, two additional algorithms from the SpliceCore platform were utilized to identify disease-specific AS changes: SpliceTrap, a method for AS quantification (Wu et al., 2011), and SpliceDuo, a regression-based predictive model for AS cross-comparison between case and control (Anczuków et al., 2015). Compared to other open access tools, SpliceCore optimizes compute time by pooling RNA-seq mapping and quantification into a single step and allows multiple cross-comparison steps to streamline AS analysis from large volumes of RNA-seq data with cloud computing (Supplemental Figure 2, Methods).

To identify productive SSO binding sites for the modulation of the AS events identified by SpliceCore, we developed a new AI/ML algorithm called SpliceLearn. Given a pre-mRNA sequence, SpliceLearn estimates the probability that delivering SSOs to a specific RNA binding position would elicit AS of its most proximal exon (Figure 1B). In addition, SpliceLearn utilizes a feature selection procedure to prioritize specific SF networks most likely to be blocked by a given SSO, increasing its usefulness and interpretability. SpliceLearn was developed as three separate AI/ML models, depending on the sequence context (i.e., exons, upstream introns, and downstream introns). Our results indicated that both intron models were far superior to the exon model (Figure 1B and Supplemental Figure 3, Methods), thus we decided to target only the upstream and downstream introns flanking exons in targetable events.

Since SpliceLearn was developed using probability-based tree learning and trained on features derived from splicing regulatory information, we were able to develop a model that is superior in both performance and interpretability. SpliceLearn outperformed comparable tools (SPANR(H. Y. Xiong et al., 2014) and MMSplice(Cheng, Yen, et al., 2019)) with more balanced sensitivity and specificity (Supplemental Figure 4). In addition, SpliceLearn performance peaked at the intuitive, optimal cutoff of a probability of 0.5, as measured by the Youden Index (Fluss et al., 2005) making SpliceLearn scores more legible to human interpretation (Supplemental Figure 4, Methods). To further the interpretability of SpliceLearn, we developed a feature selection procedure based on Shapley (SHAP) and out-of-bag (OOB) analyses (Lundberg et al., 2019). This allowed for the ranking of SpliceLearn hits by informing the most likely SFs to be blocked by SSOs at a given position on the RNA. The predicted regulatory role of these SFs was further supported by external eCLIP data from the ENCODE project (van Nostrand et al., 2020). When we examined a set of 2,124 cassette exon events, we found that the SFs determined to be most likely to regulate AS were significantly enriched for eCLIP peaks at SpliceLearn predicted binding locations (Supplemental Figure 5, Methods). Ultimately, the interpretability of SpliceLearn allows for the identification of SFs most likely to be modulated by SSOs, and therefore facilitates efficient and informed downstream mechanistic studies.

### Identification of novel AS events in TNBC using the SpliceCore platform

TNBC is one of the most aggressive forms of breast cancer (Garrido-Castro et al., 2019). Interestingly, we observed that 81.5% of TNBC samples in TCGA presented copy-number or transcriptional alterations in at least one regulatory SF, illustrating the extent of variability and potential damage to the spliceosome in TNBC (Supplemental Figure 5A). It has been shown that TNBC progression and survival depends on SFs like SRSF1 and TRA2B (Anczuków et al., 2015; Du et al., 2021; Leclair et al., 2020). Based on the unmet need and the strong scientific premise, we decided to investigate novel AS events critical for TNBC progression and develop AS-correcting SSOs for these potential targets. TNBC has been shown to be transcriptionally distinct when compared to other subtypes of breast cancer and particularly with respect to luminal breast cancer, which overexpresses hormone receptors (Kahles et al., 2018). We applied the SpliceCore platform to TCGA breast cancer data and identified 1,701 AS events expressed in TNBC basal tumors (n=169) but not normal breast tissue (n=108), and 652 AS events unique to TNBC basal tumors when compared to luminal tumors (n=694) (Figure 2A).

**Figure 2.**
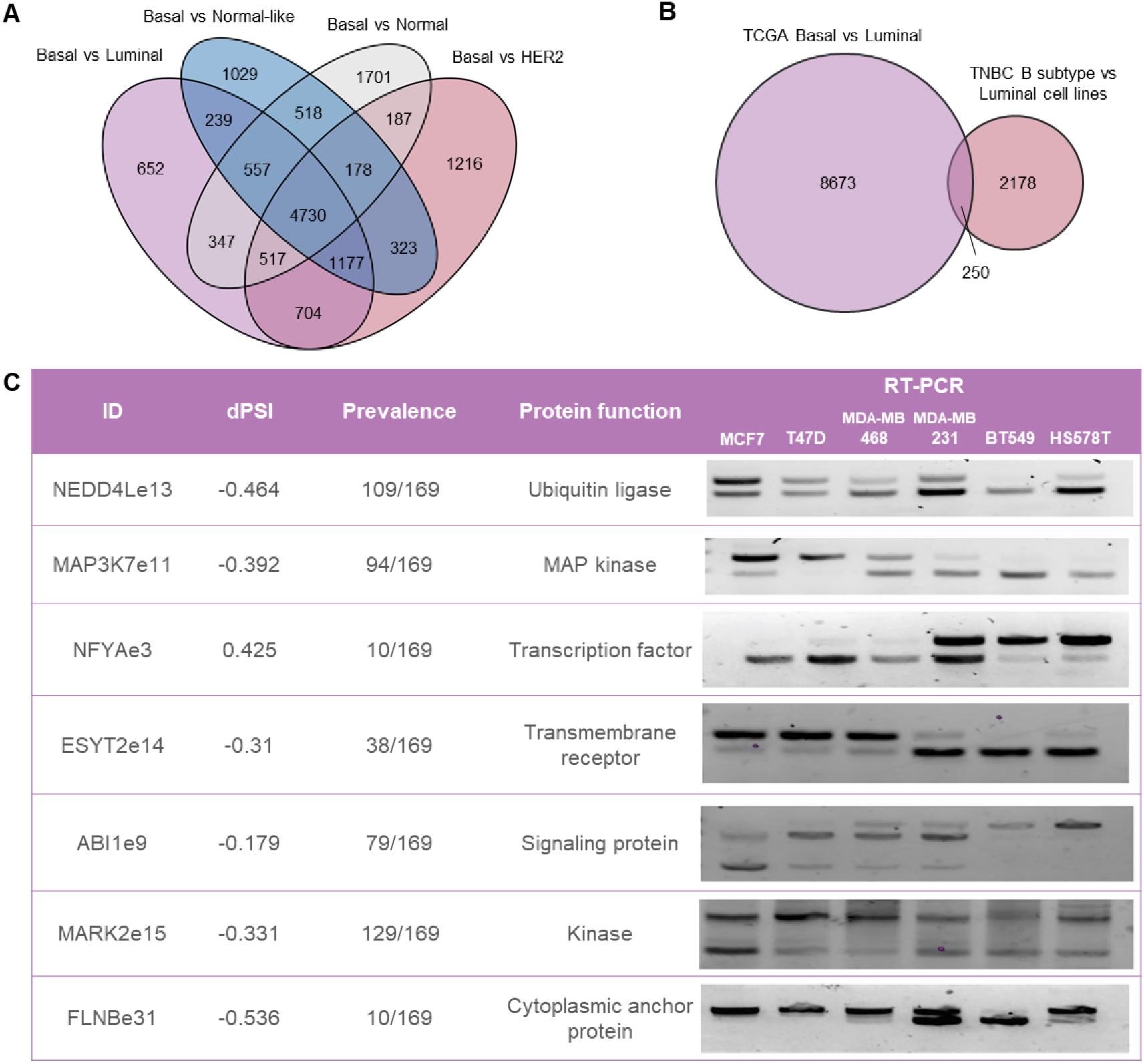
SpliceCore identified disease-specific, biologically relevant AS events in TNBC. A. Overlap of AS changes across breast cancer tissue types in TCGA. B. Overlap of AS changes identified in TCGA and in-house cell-line RNA-seq data C. Top seven SpliceCore targets identified in TNBC. The table shows dPSI values for basal vs. luminal tumor cross-comparisons, prevalence across 169 TNBC samples, function of the target candidates, and RT-PCRs with AS changes.

To independently confirm basal-specific AS changes identified in TCGA, we generated RNA-seq data in triplicate for two representative basal cell lines that emulate TNBC (HS578T and BT549) and two representative luminal cell lines as a control (MCF7 and T47D). Cross-comparison of AS profiles in basal vs. luminal cell lines resulted in the confirmation of 250 AS changes originally found in TCGA (Figure 2B). These candidates were further prioritized based on additional parameters, including the extent of AS (measured as the delta in Percent Spliced In, or dPSI), prevalence in TCGA samples, and biological significance of the underlying genes in cancer pathways. While some of SpliceCore’s candidates have been previously reported to play a role in cancer (*FLNB, MAP3K7, NFYA, ESYT2* (de Miguel et al., 2016; Dolfini et al., 2019; J. Li et al., 2018; Z. Li et al., 2021)) others were identified to be breast cancer-relevant for the first time (*NEDD4L, MARK2, ABI1*). The combination of these parameters resulted in a short-list of 7 candidates with potential for further investigation as SSO targets, and we confirmed the AS changes using RT-PCR in a panel of normal and breast cancer cell lines (Figure 2C).

### NEDD4Le13 was identified as a potential candidate for therapeutic SSOs in TNBC

AS of *NEDD4Le13* was selected by SpliceCore as a top candidate, showing prominent exon skipping in 109/169 (64%) of TNBC patients, as well as at the RNA and protein levels in breast cancer basal cell lines (Figures 2C, 3A). NEDD4L belongs to the ubiquitin ligase family of proteins that play a role in the mono-ubiquitination of several important proteins involved in cellular homeostasis and signaling, including proteins in the TGFβ pathway. At the protein level, skipping of NEDD4Le13 was predicted to remove a short loop region next to the second WW domain, a region important for binding other proteins (Figure 3B-C)(Aragón et al., 2012; Gao et al., 2009). The loop also contains an accessible threonine residue that is likely phosphorylated by Protein Kinase A.(Snyder et al., 2004) Since this region has been shown to critically interact with SMAD proteins to regulate TGFβ signaling, we hypothesized that loss of this short loop through AS of exon 13 deregulates TGFβ signaling to promote tumor progression. Accordingly, we observed that breast cancer patients expressing full-length *NEDD4L* had significantly better overall survival compared to patients with predominant skipping of exon 13 (Figure 3D). Tumor-type stratification of *NEDD4Le13* in TCGA data showed the skipping isoform to be significantly enriched in TNBC patients compared to normal breast tissue, luminal tumors, normal-like tumors, and HER2+ breast cancer samples. Of note, while differences were highly significant at the splicing level (Figure 3E), there was only a modest difference in the RNA expression of *NEDD4L* across cancer subtypes (Figure 3F), reinforcing the usefulness of AS analysis for tumor-specific target discovery.

**Figure 3.**
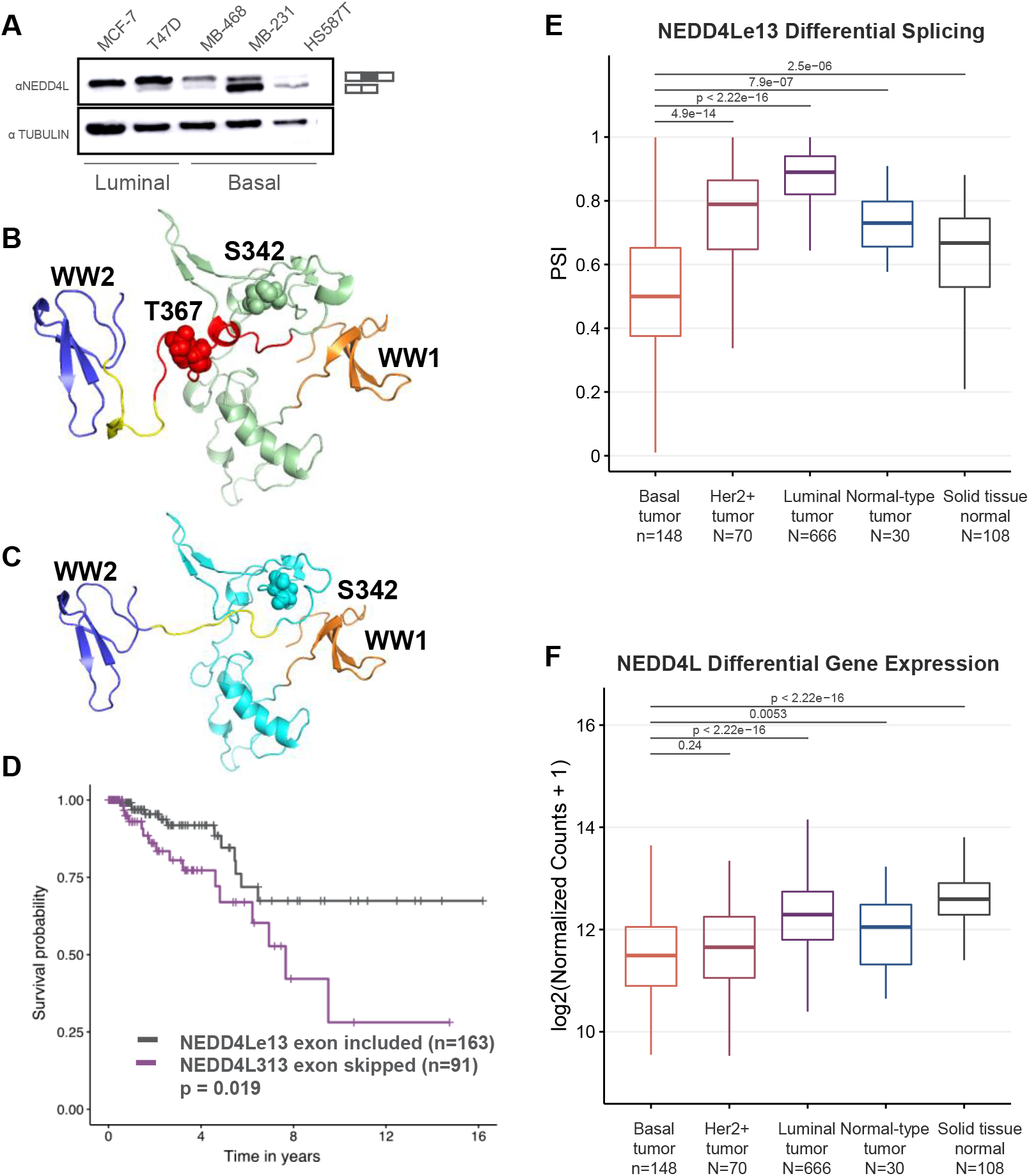
SpliceCore platform identifies NEDD4Le13 as a potential therapeutic target for TNBC. A. Western blot showing the expression of the *NEDD4L* full length (top band) and exon 13 skipping (bottom band) isoforms across a panel of breast cancer cell lines B. 3D model of amino acids 193-418 in NEDD4L from the transcript including exon 13. WW domains 1 and 2 are in orange and blue, respectively. Phosphorylation of S342 and T367 are shown as spheres. Red indicates region encoded by exon 13. Yellow colored region in both models have RMSD of 4.271. C. 3D model of the same region of NEDD4L when exon 13 is skipped. Exclusion of 20 amino acids in the loop connecting WW1 and WW2 domain alters the helix in the proximity of the WW2 domain with RMSD of 4.271 whereas no significant RMSD (0.188) was obtained when the two full models were superimposed. D. Survival curves showing a significant difference in survival between patients where *NEDD4Le13* is included and *NEDD4Le13* is skipped. D. *NEDD4L* Percent Spliced In (PSI) and E. gene expression data from the TCGA BRCA samples stratified into subtypes.

### SpliceLearn identified an optimal SSO to promote *NEDD4Le13* inclusion and explained the underlying AS regulatory network

Next, we utilized SpliceLearn to find the optimal binding sites for SSOs to promote *NEDD4Le13* inclusion, as well as to identify the underlying SF network regulating this AS event. SpliceLearn analysis revealed that the three SFs most likely to bind the upstream intron and regulate AS of *NEDD4Le13* were HNRNPL, QKI, and SRSF7.

These three SFs showed confirmatory ENCODE eCLIP peaks (van Nostrand et al., 2020) overlapping with SpliceLearn hits. Particularly, HNRNPL presented a strong binding signal in four eCLIP replicates, in stark contrast to other well-known SFs like SRSF1, which SpliceLearn did not predict to regulate *NEDD4Le13* splicing, and did not show any confirmatory eCLIP peaks (Figure 4A). We found that *HNRNPL* and *SRSF7* were both significantly overexpressed in basal tumors when compared to other breast cancer subtypes and to normal tissue samples (Figure 4B-C). QKI was overexpressed in TNBC when compared to luminal and HER2+ samples but was downregulated when compared to normal-like tumors and normal tissue (Figure 4D). These subtype-specific changes in SF expression highlight their potential to regulate tumor-specific AS events like *NEDD4Le13*. Finally, we investigated protein-protein interactions (PPIs) among the three SFs using a probabilistic network of spliceosomal PPIs (Akerman et al., 2015). In fact, we observed that while QKI and SRS7 were unlikely to bind with each other directly (*P*_*QKI−SRSF7*_ = 0.098) the probability of each directly binding HNRNPL was high (*P*_*HNRNPL−QKI*_ = 0.826, *P*_*HNRNPL−SRSF7*_ = 0.784) (Figure 4E). Of note, PPIs predicted between HNRNPL and QKI/SRSF7 were among the top interactions within the full network of HNRNPL interactions that included 601 proteins (Figure 4F). In summary, these data suggest that HNRNPL binds consistently and specifically to the intron upstream of *NEDD4Le13*, to further recruit QKI and SRSF7 as they work together to regulate AS.

**Figure 4.**
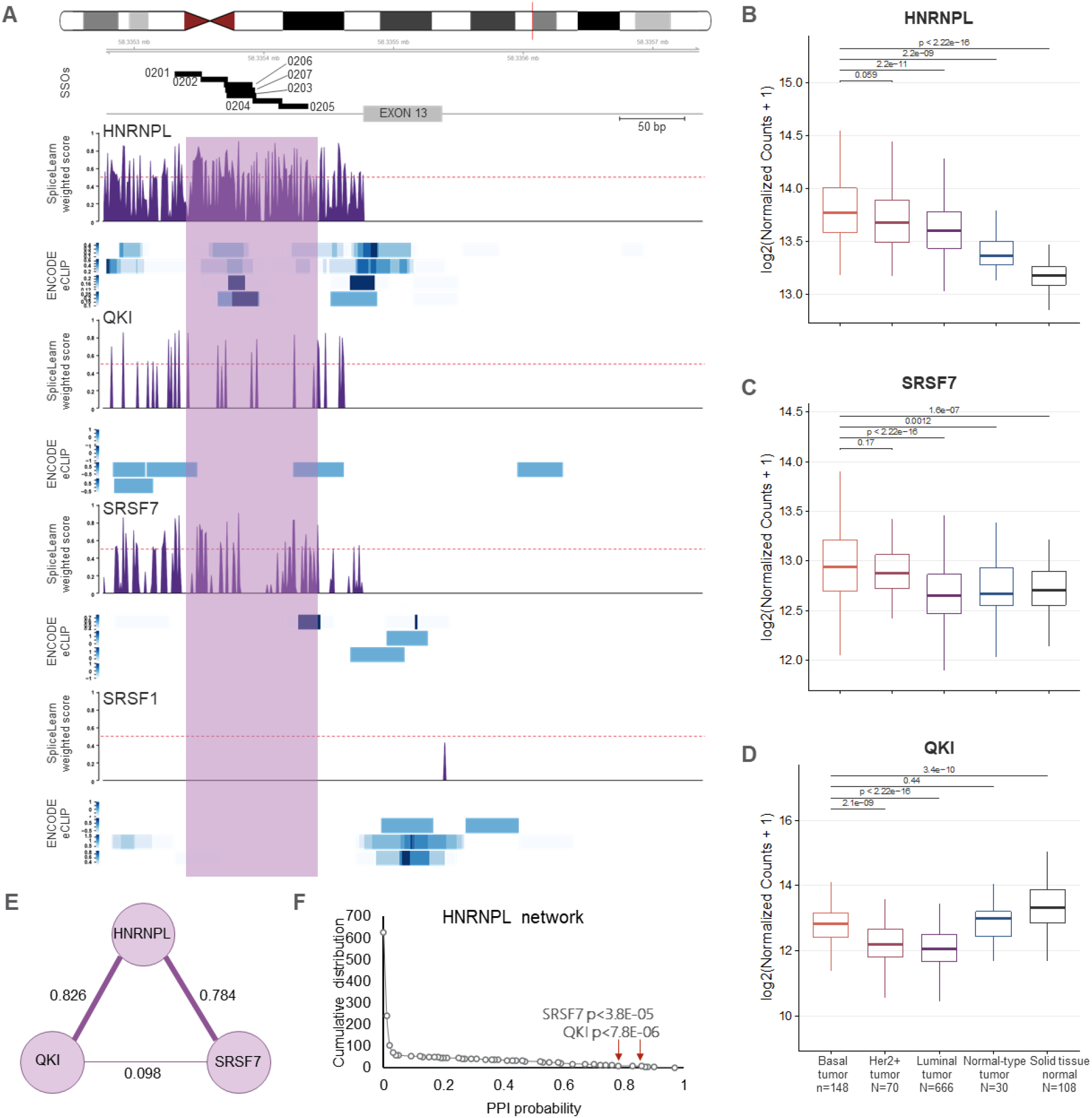
SpliceLearn predicts the optimal functional SSO binding site to modulate the splicing of NEDD4Le13 in TNBC. A. SF-specific SpliceLearn scores as determined by SHAP analysis. SRSF1 was included as a negative example. Black boxes in the top track indicate the binding sites of the SSOs (20-mers) designed and tested to promote *NEDD4Le13* exon inclusion. Blue heatmaps indicate eCLIP binding from ENCODE cell lines, where darker blue indicates higher binding score. B-D. *HNRNPL, SRSF7*, and *QKI* expression levels across the subtypes of breast cancer in TCGA. E. Protein-protein interaction network between the three SFs highlighted in A as determined in Akerman et al 2015. F. Probability distribution of protein-protein interaction scores between *HNRNPL* and the rest of the SFs in the Akerman et al 2015 dataset. P-values and probability highlighted for *QKI* and *SRSF7*.

To experimentally validate the SpliceLearn-predicted functional SSO binding sites, a list of 20- to 22-mer sequences spanning locations across the highest SpliceLearn scores that overlapped with ENCODE HNRNPL peaks was generated (Figure 4A, Table S1). These seven oligos were chemically modified to enhance stability and nuclease resistance using 2^nd^ generation antisense chemistry consisting of a phosphothioate backbone and uniformly modified 2’ methoxyethane (2’MOE) ribose sugar. Chemically synthesized and purified oligonucleotides were subjected to functional assays in breast cancer cell lines. Out of the seven sequences tested, SSO-0205 was found to promote the strongest *NEDD4Le13* inclusion in MDA-MB-231 cells. Treatment with SSO-0205 caused an average of 67% exon inclusion in 3 independent experiments, compared to 15% inclusion in the lipofectamine control group (Figure 5A-B). Additionally, MDA-MB-231 cells treated with SSO-0205 showed substantial exon 13 RNA-seq read coverage in *NEDD4L*, compared to the untreated, lipofectamine treated, or SSO-0202 treated cells (Figure 5C). In summary, SpliceLearn successfully identified functional SSO binding sites, reducing the need for time-consuming microwalks for SSO optimization.

**Figure 5.**
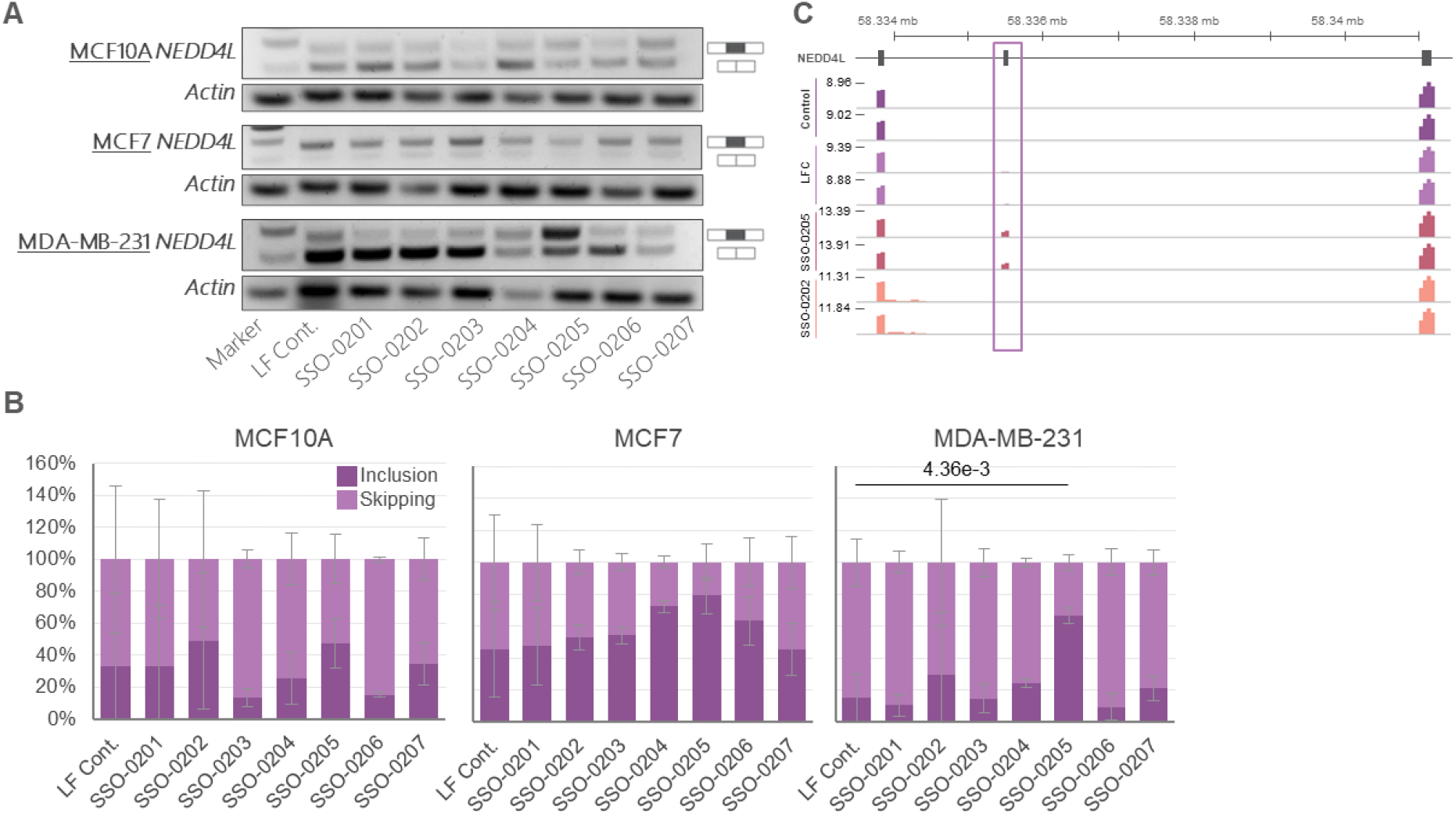
SpliceLearn-predicted SSOs modulate *NEDD4Le13* inclusion. A. Representative 2% agarose gel for *NEDD4L* PCR showing isoform changes in three breast tissue cell lines treated with the indicated SSOs (400nM) for 48h. Actin expression used as cDNA internal control. B. Inclusion/Skipping percentage measured by qPCR in three breast cancer cell lines after SSO (400nM) treatment for 48h (n=3). C. Genome browser tracks displaying RNA-seq data in MDA-MB-231 cells (untreated (dark purple), treated with a lipofectamine control (light purple), SSO-0205 (maroon), or SSO-0202 (peach). Box highlights *NEDD4Le13*.

### SSOs targeted to NEDD4Le13 promoted exon inclusion and affected the TGFβ pathway

NEDD4L is a ubiquitin ligase that has been previously shown to play a role in TGFβ regulation (Aragón et al., 2012; Gao et al., 2009). Moreover, NEDD4L has been shown to be involved in both oncogenesis and tumor suppression (X. Y. Guo et al., 2022; Xie et al., 2021). However, it is unclear if a specific AS isoform of NEDD4L is responsible for regulating TGFβ signaling in TNBC tumors, where it has the potential to drive multiple aspects of tumor progression. The TGFβ-dependent response is highly contextual throughout development, across different tissues, and therefore its dysregulation is highly relevant in tumor development and progression (Bellomo et al., 2016; Massagué, 2008).

Using MDA-MB-231 as a representative TNBC cell line, and MCF10A (cell line derived from breast fibroadenoma) as a control, we evaluated the splicing patterns of *NEDD4Le13* in the context of TGFβ stimulation via treatment with human recombinant TGFβ (hrTGFβ). Untreated MDA-MB-231 cells predominantly expressed the skipping isoform. When SSO-0205 was added to the cells for 24 hours, there was a significant increase in *NEDD4Le13* inclusion. SSO-0205 maintained a high level of *NEDD4Le13* inclusion even after exposure to hrTGFβ for 3 or 6 hours. On the other hand, even though MCF10A cells often skip *NEDD4Le13* in untreated conditions, they did not show a response to SSO-0205 under TGFβ treatment, perhaps due to differences in *NEDD4Le13* regulation between the cell types and a dependency on the short isoform in the context of tumor maintenance (Figure 6A). Accordingly, RNA-seq analysis showed *HNRNPL, SRSF7*, and *QKI* to be highly expressed in MDA-MB-231 cells compared to MCF10A cells, pointing to a specific role of this regulatory network in the context of TNBC (Figure 6B).

**Figure 6.**
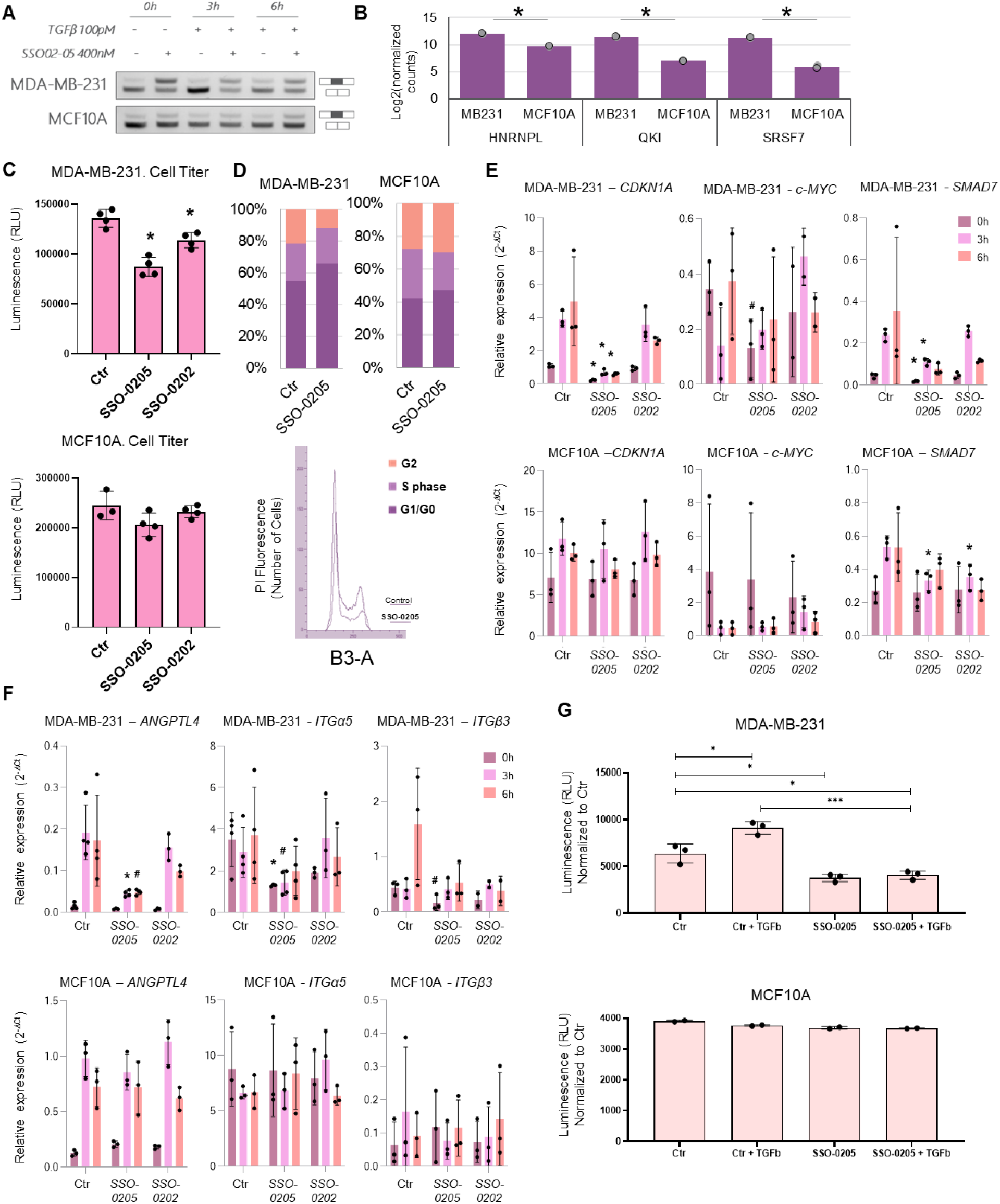
SSO-0205 modulates *NEDD4Le13* inclusion, causes cancer cell migratory response to TGFβ, and decreases cell proliferation. A. PCR measuring *NEDD4L* isoforms in MDA-MB-231 and MCF10A cells in response to TGFβ stimulation (0, 3 or 6 hours) after SSO-0205 treatment (400 nM) (24h). B. Normalized RNA-seq counts for HNRNPL, QKI, and SRSF7 in MCF10A and MDA-MB-231 cells showing significantly higher expression of these SFs in MDA-MB-231 cells. Statistical differences calculated with DESeq2; *≤1×10^−85^ (n=2). C. Cell proliferation as measured by Cell TiterGlo® 96 well assay. MDA-MB-231 (top graph) and MCF10A (bottom graph) cells were treated with lipofectamine alone as a control or with the indicated SSO for 48h (400 nM) (n=4). D. Cell cycle phase analysis measured by propidium iodide flow cytometry. Cell cycle phase for each cell line is represented by % (top graphs), a representative Propidium Iodide histogram plot for MDA-MB-231 is shown below. Cells were treated with lipofectamine alone or with SSO-0205 (400 nM) for 24h (n=3). E. qPCR quantifying expression levels of the indicated cell cycle related genes in MDA-MB-231 (top line) or MCF10A (bottom line) cells treated with SSO or lipofectamine alone as control for 24 hours followed by 0, 3 or 6 hours treatment with hrTGFβ (n=3-4). F. qPCR quantifying expression levels of the indicated migration/invasion related genes in MDA-MB-231 (top line) or MCF10A (bottom line) cells treated with SSO (400 nM) or lipofectamine as control for 24 hours followed by 0, 3 or 6 hours treatment with hrTGFβ (n=3-4). G. Transwell migration assay on the indicated cell types (MDA-MB-231 top graph and MCF10A bottom graph), cells were treated with lipofectamine or SSO-0205 (1 μM) for 24 hours, followed by overnight TGFβ treatment in the Transwell chamber (n=2-3). Statistical differences were calculated by Student’s t-test vs the corresponding time point at the control (Ctr) group; *≤0.05; #≤0.1.

Next, we evaluated the effect of SSO-0205-induced *NEDD4Le13* inclusion on the protein level and localization of several members of TGFβ pathway. MDA-MB-231 cells were treated with either SSO-0205, SSO-0202 (used as a control), or with lipofectamine alone as a vehicle control for 24 hours followed by a 0-, 1- or 3-hour treatment with hrTGFβ. Following SSO-0205 and hrTGFβ treatment, MDA-MB-231 cells showed an accumulation of phosphorylated SMAD2/3 (phSMAD2/3) and TGFβ-receptor (TGFβRI) in the cytoplasm, concomitant with a nuclear decrease in phSMAD2/3, total SMAD2/3 and a decrease in TGFβRI at the membrane compared to the lipofectamine control (LFC). These cellular localization changes were time dependent and more prominent at 3 hours after the hrTGFβ treatment. In the case of SSO-0202 and in MCF10A cells, these effects on protein localization were very modest, as expected given the <20% change in inclusion ratio in cells treated with SSO-0202 and the lack of response to the SSOs by MCF10A cells (Supplemental Figure 7).

NEDD4L has been previously described to interact with phSMAD2/3, SMAD7 and TGFβRI as part of its role in the ubiquitination and subsequent proteasomal degradation of these proteins (Gao et al., 2009; Kuratomi et al., 2005). We used MMG132 to inhibit proteasome activity and performed immunoprecipitation assays for SMAD2/3 and TGFβRI to determine their interaction with NEDD4L after treating cells with SSO-0205 for 24h then with hrTGFβ for 0, 3 or 6 hours. When we inhibited proteasome activity before treating with SSO-0205 the total levels of NEDD4L, phSMAD2/3 and TGFβRI increased after hrTGFβ addition to the cells at 3 and 6 hours, specifically in the MDA-MB-231 cells (Supplemental Figure 8A). Moreover, we found that SSO-0205 treatment promoted PPI between NEDD4L and SMAD2/3 as well as between NEDD4L and SMAD7, particularly after 3 hours of hrTGFβ treatment as observed by an increase of NEDD4L signal after SMAD2/3 IP and SMAD7 IP (Supplemental Figure 8A). Importantly, MCF10A cells treated with SSO-0205 and hrTGFβ did not show any change in downstream TGFβ signaling protein subcellular localization or PPIs by IP (Supplemental Figure 8A). These subcellular fractionation data and PPI results confirm the direct role of the NEDD4L inclusion isoform in modulating TGFβ pathway activity by marking the phSAMDs and TGFβRI for degradation as previously described (Gao et al., 2009; Kuratomi et al., 2005).

### SSO-mediated NEDD4L exon inclusion selectively modulated TNBC cell proliferation

To understand the role of the NEDD4Le13 in TNBC cell viability, we conducted a cell titer glow viability assay in the presence or absence of SSO-0205. Treatment of MDA-MB-231 cells with SSO-0205 significantly decreased the viable cell count in MDA-MB-231 cells both in fixed concentration (400nM), as well as in a dose-dependent manner. (Figure 6C, Supplemental Figure 8B). Additionally, a cell cycle evaluation indicated that SSO-0205 treatment significantly increased the number of cells in G1 and decreased the number of cells in G2 at the 24-hour time point (Figure 6D). Treatment of MCF10A with SSO-0205 did not change cell proportions along the cell cycle or cause a significant change in cell viability. To determine whether NEDD4Le13 AS modulates the cell cycle in response to TGFβ, we analyzed the gene expression of *CDKN1A* and *c-MYC*, two key downstream targets of TGFβ in epithelial cells (Decker et al., 2021; Weiss, 2003). We observed that 24 hours after SSO-0205 treatment, and prior to hrTGFβ treatment, MDA-MB-231 cells showed a decrease in both *CDKN1A* and *c-MYC* gene expression levels compared to the control cells treated with lipofectamine (Figure 6E). Interestingly, when hrTGFβ was added to the cells, the increase in *CDKN1A* gene expression observed in the lipofectamine control group was inhibited by the SSO-0205 treatment (Figure 6E) (TGFβ time 3 and 6h). Additionally, we saw that *SMAD7* expression showed a similar expression profile to the *CDKN1A* when comparing the lipofectamine control to SSO-0205 treated cells (Figure 6E). Consistent with the viability assay, these changes were limited to MDA-MB-231 cells and not observed in MCF10A cells. Taken together, these data indicate that promoting *NEDD4Le13* inclusion with SSO-0205 specifically modulated the proliferation and viability of the MDA-MB-231 cells, further supporting its biological importance and therapeutic potential in TNBC.

### NEDD4Le13 AS modulated the MDA-MB-231 migratory response to TGFβ pathway activation

Cell migration and invasion is one of the hallmarks of cancer that enables metastasis. *ANGPTL4, ITGα5*, and *ITGβ3* are TGFβ-dependent cell migration-associated markers. SSO-0205 pre-treatment of MDA-MB-231 cells decreased the gene expression response of *ANGPTL4, ITGα5*, and *ITGβ3* after hrTGFβ addition at time 0, 3 and 6h, while SSO-0202 treatment did not cause significant changes for any of the time points (Figure 6F). Expression changes in migration-associated genes were not observed in MCF10A cells after SSO-0205 or SSO-0202 treatment, even though TGFβ stimulation induced the expression of some of these genes (Figure 6F). Finally, evaluation of changes in the migration of MDA-MB-231 cells using a Transwell Migration Assay showed a 40-50% decrease in cell migration in SSO-0205 treated cells, even in the presence of hrTGFβ. This effect was specific to MDA-MB-231 cells; MCF10A cells did not show a decrease in migration in the presence of SSO-0205 (Figure 6G).

Overall, we have shown that NEDD4Le13 skipping contributes to TNBC tumor progression and promotes overactivation of the TGFβ pathway in MDA-MB-231 cells. Furthermore, SSO-0205 was able to reverse *NEDD4Le13* exon skipping by presumably blocking the binding of HNRNPL, QKI and SRSF7 to the intron upstream *NEDD4Le13*, thereby decreasing both the proliferative and migratory behavior of MDA-MB-231 cells. Both the target, *NEDD4Le13*, and the binding site for the compound, SSO-0205, were discovered *de-novo* by applying AI/ML algorithms to RNA-seq data sets, demonstrating the potential for innovative drug discovery with the SpliceCore platform in the field of RNA therapeutics.

## Discussion

Our goals for this study were two-fold: to develop a new SSO compound to modulate AS of a novel drug target, and to do so through a data-driven approach, using RNA-seq data and AI/ML algorithms. Here, we experimentally validated *NEDD4Le13*, a new target for TNBC, along with a corresponding SSO called SSO-0205. Both were predicted *de-novo* using the SpliceCore platform. SpliceCore allows for drug discovery at scale because it has been optimized for AS analysis of large RNA-seq datasets, outperforming many of the most common open-access tools (Supplemental Figure 2). SpliceCore’s ability to rapidly integrate and analyze thousands of RNA-seq samples to identify druggable AS events reinforces its competency as a target-discovery platform for SSO development.

The binding site for the SSO-0205 compound that successfully promoted *NEDD4Le13* inclusion was predicted *de-novo* using SpliceLearn. In terms of interpretability, SpliceLearn outperforms comparable methods (Supplemental Figure 4). We have improved the interpretability of SpliceLearn by employing a novel feature selection method using SHAP/OOB. Previous studies showed that while many predictive models in molecular biology are optimized for maximum performance, they often lack interpretability, making it harder for scientists to make actionable decisions based on the AI/ML algorithm’s results. Reduced interpretability can stem from the use of complex predictive features with hidden associations, or unbalanced sensitivity/specificity, often overlooked in favor of maximizing performance metrics such as AUC (Azodi et al., 2020; Johansson et al., 2011). In contrast, SpliceLearn relies on the use of tree-based learning methods, which are inherently more interpretable (e.g., when compared to deep learning) because they enable the investigation of the content and relationship between predictive features. SpliceLearn therefore allows for the identification of regulatory SFs at each nucleotide of interest, which is crucial for downstream studies on the regulation of the AS event and the mechanisms by which the SSOs modulate it (Figure 4, Supplemental Figure 5). SpliceLearn optimizes both sensitivity and specificity, which we found to be lacking in comparable algorithms (Supplemental Figure 4). Finally, SpliceLearn eliminates the need for time consuming and expensive SSO microwalks that require 100-200 tiling oligos to be tested manually for activity. ML-based SSO design not only consolidates the number of oligos tested, but also informs the potential regulatory mechanism associated with the targeting sites.

Remarkably, the SpliceCore platform validation study presented here produced significant biological insights related to splicing regulation in TNBC. First, we identified seven AS events highly recurrent in TNBC. Of particular interest, AS of *NEDD4Le13* was found to be present in 64% of TNBC patient tumors. Second, we showed that NEDD4Le13 is a promising target for TNBC since splice-switching with SSO-0205 caused TNBC-specific viability loss (Figure 6). This effect may be partially explained through altered localization of SMAD proteins downstream of pro-tumorigenic TGFβ activation (Supplemental Figure 7). The tumor-specific anti-proliferative effects of SSO-0205 are encouraging and may result in reduced toxicity compared to chemotherapy in future pre-clinical and clinical development. Third, skipping of NEDD4Le13 causes the exclusion of a protein loop region between the WW1 and WW2 domains important for protein interaction (Figure 3B-C) (Aragón et al., 2012; Gao et al., 2009). This loop region contained a threonine residue that can be phosphorylated, potentially adding an additional regulation through post translational modifications (Snyder et al., 2004). Fourth, SSO-0205 treatment affected TGFβ-dependent gene expression changes in proliferation-(*CDKN1A, C-MYC*) and invasion-related genes (*Integrins α5 and β3, ANGPTL4*), leading to significant changes in cell cycle and cell migration (Figure 6). While the TGFβ-independent downregulation of *CDKN1A* in the presence of SSO0205 was surprising, it is possible that additional players regulate *CDKN1A* levels through non-canonical TGFβ regulation (Weiss, 2003). Finally, despite the abundance of *NEDD4Le13* skipping in MCF10A cells (a non-cancerous mammary fibroadenoma cell line), SSO-0205 did not promote exon inclusion in these cells, suggesting that the TNBC-specific SF network identified with SpliceLearn could have a differentiating role, further supported by the observed TNBC-specific overexpression of *HNRNPL, SRSF7* and *QKI1*. We suggest that *NEDD4Le13* skipping is lineage-restricted to basal cells like MCF10A and MDA-MB-231; however, AS regulation appears to divert between both cell types, such that exon skipping in the latter promotes tumor progression, which may lead to dependency for *NEDD4Le13* skipping in the context of TNBC (Figure 6B).

Currently, several SSOs remain under investigation as potential treatments for cancer and other diseases, providing an alternative to small molecules, which are currently limited to a subset of “druggable” proteins.^3,10^ There is still progress to be made in the field of SSO therapeutics, especially in the area of drug delivery, but this is a highly active area of research and is the focus of several biotechnology companies (Roberts et al., 2020). Ultimately, this study illustrates the usefulness of a new AI/ML algorithm developed upon key principles of RNA biology, to extract novel, actionable insights from RNA-seq data. We have shown that it is possible to identify a novel drug target, and design an effective SSO against it, that not only modulates exon inclusion but ay lso displays anti-cancer activity. The efficiency of the SpliceCore platform in analyzing thousands of RNA-seq samples to identify novel drug targets in a timely manner can be expanded beyond cancer research into other diseases driven by splicing alterations, like neurodegeneration or metabolic diseases (Centa et al., 2020; Finkel et al., 2017; Han et al., 2020; J. Kim et al., 2019; Syed, 2016; K. R. Wagner et al., 2021).

## Methods

### SpliceCore

The SpliceCore software platform is Envisagenics’ proprietary technology for the discovery of splicing-modulatory drug targets using RNA-seq data. SpliceCore utilizes scalable cloud-computing through Microsoft Azure services to efficiently perform *de-novo* transcript assembly using an exon-centric reference transcriptome called TXdb, followed by AS analysis. SpliceCore implements the following algorithms: SpliceTrap, a Bayesian-based method for RNA-seq alignment and AS quantification, SpliceDuo, a regression-based predictive model for AS cross-comparison, SpliceImpact, an AI/ML algorithm for the prioritization of biologically relevant AS events based on the impact of AS on protein structure and RNA stability, and SpliceLearn, its newest algorithm for the identification of productive SSO binding sites (Anczuków et al., 2015; Wu et al., 2011).

### SpliceLearn

SpliceLearn, an AI/ML module of the SpliceCore platform, was trained with predictive features combining three sources of splicing regulatory information: the relative position of prospect SSOs binding sites to an exon, the identity and binding motif scores of SFs potentially blocked by SSOs (Paz et al., 2014; Ray et al., 2009; X. Wang et al., 2011), and the interaction of such SFs with other SFs within the spliceosome (Figure 1B, Akerman et al., 2015).

To train and test SpliceLearn, we utilized data from a massively parallel splicing minigene reporter assays (MFASS(Cheung et al., 2019)) which quantifies the effect of SNVs on AS outcomes, therefore mimicking the impact of SF-blocking by SSOs. The MFASS datasets assessed the effect of 27,733 SNVs extracted from the ExAC database on the splicing of 2,198 distinct human exons. Of these, 14,130 SNVs occurred in exons, 7,938 in the intronic region upstream the 3’ splice site, and 6,271 in the intronic region downstream the 5’ splice site. These three RNA regions present striking differences in sequence composition and have evolved to play distinct roles in splicing regulation. Therefore, we developed three independent SpliceLearn models SpliceLearn-e, -up and -down to account for the unique regulatory properties of each exonic and upstream/downstream intronic elements (Figure 1B and Supplemental Figure 3, Methods). To test SpliceLearn with an independent data source, we used the Vex-seq dataset, which consists of 1226 qualifying variants that were experimentally identified to have impact on pre-mRNA splicing using a high-throughput reporter assay.(Adamson et al., 2018) The SpliceLearn-down model classified VexSeq data with an AUC of 0.98, SpliceLearn-up with and AUC of 0.86 and SpliceLearn-e with AUC of 0.66 (Supplemental Figure 3B). These results confirm the usefulness of both intronic SpliceLearn models to predict SF-binding perturbation positions critical for AS regulation and useful for SSO targeting. See Methods for detailed description of SpliceLearn training, testing, and benchmarking.

### NEDD4L protein structure modelling

The three-dimensional model of the NEDD4L amino acids 193-418 was obtained by *ab-initio* modelling on Robetta platform (http://robetta.bakerlab.org/, D. E. Kim et al., 2004). Five models were obtained and all of them were validated in the Molprobity server (http://molprobity.biochem.duke.edu/, Davis et al., 2007). Two energy minimization processes using the Chimera program (Yang et al., 2012) were followed by residue specific minimization in SPDBV (Guex & Peitsch, 1997). The model with best molprobity score was used for comparative modelling of alternate structure with exclusion. Root mean square deviation was calculated by superimposing two structures in PYMOL.

### Cell Lines

All cell lines used in this study were purchased from NCI Cancer Cell line-60 panel and/or American Type Culture Collection (ATCC) authenticated by STR profiling and mycoplasma testing.

### RNA extraction and RT-PCR

Total RNA from different breast cancer cell lines were extracted using the TriZol reagent as per manufacturer’s instruction. One microgram of RNA was used to make cDNAs using oligo-dT primers. One-tenth of the volume of the cDNA reaction mixture was used in the PCR reaction containing specific forward and reverse primers for detecting different splicing isoforms. The amplicons are separated in 2% agarose gels. The primer sequences for genes used in the PCR assays are given below.

### Splice-switching assays

Computationally predicted SSO sequences were purchased from Microsynth technologies, Switzerland. The oligos were uniformly modified to contain phosphothioate backbone and 2’ methoxy ethane containing ribose sugars (2’ MOE). All oligos were HPLC purified and lyophilized oligos were reconstituted in calcium and magnesium free PBS to obtain 100uM stocks. The cells were plated to 80% confluency and 400nM SSO specific to *NEDD4L* or control SSOs was transfected using Lipofectamine 3000 according to the manufacturer’s instruction. Alternatively linear concentration of doses of the oligos were used for the dose response assay. The cells were harvested, and RNA was extracted 48 hours post-transfection. RT-PCR evaluation of the splice-switch was performed as described above.

### Cell viability assay

Cell TitreGlo® (Promega) was used to determine the viability of the SSO transfected breast cancer cells according to manufacturer’s instruction. Briefly, about 10K MCF7 or MDA-MB-231 cells were plated into each well of a 96 well plate that was treated with either control or single concentration or linear concentrations (100nM, 200nM, 400nM, 600nM, 800nM, 1uM) of *NEDD4L* specific SSOs in the presence of lipofectamine in triplicates. 48 hours post-transfection equal volume of Cell TitreGlo reagent was added to the well and the total luminescence was measured using a plate reader (Perkin Elmer). The average percentage of viable cells that is proportional to the total luminescence was calculated and plotted to obtain viable cell percentage and dose response curve for *NEDD4L* SSO.

### Cell culture and cell treatments for TGFβ pathway studies

MDA-MB-231 cells were grown in RPMI media containing 10% fetal bovine serum (FBS) and penicillin and streptomycin. MCF10A cells were grown in DMEM/F12 media supplemented with MEGM™ Mammary Epithelial Cell Growth Medium SingleQuots™ Kit (Lonza, CC-4136) and penicillin and streptomycin. For the TGFβ pathway analysis experiments, cells were plated in 6 well plates at 2×105 cells/well. SSOs treatment were added to the cells growing in complete media at 400nM final concentration, in the presence of lipofectamine 3000 (Thermo Fisher Scientific, L3000001) following manufacturer instructions, unless otherwise specified. Lipofectamine alone was added as control group. Human recombinant TGFβ (hrTGFβ) (R&D Systems), was administered directly into the media at 100pM final concentration. RNA or protein samples were harvested at different time points. Nuclear, Cytoplasm and membrane protein fractions were extracted following the manufacturer instruction for the Subcellular Protein Fractionation Kit for Culture Cells (Thermo Scientific, Cat. # 78840).

### Cell Cycle Analysis

MDA-MB-231 and MCF10A cells were plated on 6 well plates at 1×105 cells/mL. The following day the cells were treated with SSO-0205 at 400nM for 24 hours. Cells were trypsinized, fixed with 70% ethanol on ice and stained with Propidium Iodide (Invitrogen, P3566) at 50μg/mL final concentration for 30-40 minutes. Stained cells were analyzed by flow cytometry MACSQuant® Analyzer (MACS Miltenyi Biotech) following the manufacturer instruction.

### Transwell migration assay

MDA-MB-231 and MCF10A cells were plated on 6 well plates at 1×105 cells/mL, treated with SSO-0205 at 1μM (no lipofectamine – free uptake) for 24 hours, trypsinized and resuspended in serum free media at 5×105 cells/mL containing SSO-0205 at 1μM with or without hrTGFβ 100pM. 500μL of cell suspension was added to the transwell inner chamber (8μm pore size) (Falcon, #353097), the inserts were placed on a 12 well plate containing complete media to promote chemoattraction to the outer chamber. Cells were incubated overnight and migratory cells were measured following the manufacturer instruction for the cell dissociation and Calcein AM staining and luminescence measurement (Cultrex® Cell Invasion Assay, Trevigen, Cat. # 3455-024-K).

### RNA-sequencing

∼5*105 cells were harvested in biological triplicates from each breast cancer cell line (MCF7, T47D, MDA-MB-231, MDA-MB-468, HS578T, BT549). Frozen cell pellets were provided to GeneWiz (SD, California) for RNA extraction and library preparation. Briefly, 1ug of RNA extracted using TriZol was enriched for PolyA containing RNA using oligo dT columns. Libraries for polyA+ RNA-seq were prepared using TruSeq chemistry (Illumina), multiplexed, and sequenced to obtain paired-end 101-base-pair (bp) reads on an Illumina HiSeq 2000 platform, resulting in 30 million to 45 million reads per library.

## Supplemental Methods

### Assembly of TXdb

TXdb is an exon-centric reference transcriptome used by all of SpliceCore’s algorithms (Wu et al., 2011). The premise of “exon-centric” is to treat the transcriptome as a collection of independent AS events rather than full-length transcripts. In TXdb, CA and IR events are represented as exon trios where the middle exon is subjected to AS analysis and the flanking exons provide the transcriptomic context with the corresponding splicing junctions necessary for the analysis. Every exon trio is presented in two splicing states: inclusion, where the three exons are connected through a pair of splicing junctions, and skipping, where the flanking exons are connected by a single splicing junction and the middle exon is skipped. Likewise, alternative 3’ splice sites (A3SS) and alternative 5’ splice sites (A5SS) are represented as exon duos where the extended exon segment is subjected to AS analysis and the remining sequences provide context (Supplemental Figure 1A). For simplicity, we use the term “inclusion” to also define intron retention and distal A3SS and A5SS splicing; and we use “skipping” to define intron splicing and proximal A3SS and A5SS splicing. In addition, CA events in TXdb include annotations for both alternative and constitutive exons, i.e., for exon trios only supported by their inclusion state. In contrast, evidence for both inclusion and skipping states were required for IR, A3SS and A5SS as a way to limit the sequence search space due to lengthy introns and the exponentially large number of dinucleotides matching the splice site consensus that could result in A3SS or A5SS. To generate TXdb, we assembled mRNA contigs from 1,252 breast cancer RNA-seq datafiles from TCGA. STAR aligner (Dobin et al., 2013) was used for read mapping and Stringtie (Pertea et al., 2015) for contig building. A total of 5,617,407 AS events were supported by at least one out of 1,252 RNA-seq datafiles, although many of these AS events were redundant. Almost every AS was clustered with several others showing identical inclusion and skipping junctions, identical middle (or extended) exon, but flanking exons of different sizes. We pruned redundant clusters of AS events by selecting a single representative exon trio/duo by applying the following steps: First, we looked for known AS events supported by Ensemble or RefSeq. Second, we prioritized exon trios/duos supported by Ensemble or RefSeq. Third, we took AS events with the highest reliability score (see next section for details). Finally, if a tie persisted, we selected the shortest AS event to further reduce sequence search space (Supplemental Figure 1B). We identified a total of 1,743,426 non-redundant AS events, including 1,190,514 CA, 199,238 IR, 202,851 A3SS and 150,823 A5SS (Supplemental Figure 1C). Also 534,231 (31%) exon trios/duos were supported by ENSEMBL or RefSeq (GRCh38.p12), 77,381 (4%) further presented both inclusion and skipping evidence in ENSEMBL or RefSeq (i.e. known AS events). The remaining 1,131,814 (65%) showed no evidence of in public mRNA databases and where therefore annotated as novel trios/duos (Supplemental Figure 1D).

### SpliceCore analysis and benchmarks

SpliceCore takes RNA-seq FASTQ/A files as an input to predict disease-specific SSO drug targets. The first step of the analysis is to estimate AS between “disease” and “normal” RNA-seq samples in order to prioritize drug targets amenable to SSO modulation. AS analysis is often divided into three steps: alignment, quantification, and comparison (Alamancos et al., 2014) (Supplemental Figure 2A). The SpliceCore platform uses the SpliceTrap algorithm to align RNA-seq data to TXdb and quantify the “percent spliced in” (PSI) of every AS event. Next, SpliceDuo performs case/control comparisons and reports splicing changes as ΔPSI values between -100% (i.e. full exon skipping) and 100% (full exon inclusion). Sequence alignment is the most time-consuming step of RNA-seq analysis, in part due to the use of a large reference transcriptome like TXdb, with 1,743,426 AS annotations. However, most RNA-seq analysis projects only require a single alignment iteration. One-time sequence alignment is common practice in bioinformatics supported by evidence that changes in alignment parameters have little impact on both technical and biological performance.(Ballouz et al., 2018) The SpliceTrap algorithm is optimized for a one-time execution step that includes sequence alignment and PSI quantification. In contrast, the comparison step is often repeated multiple times to allow thorough interrogation of the data (Supplemental Figure 2A). This is especially important when analyzing RNA-seq data from heterogenous patient cohorts, including subjects with various disease subtypes, at different disease stages, responding differently to drug treatments, and with diverse clinical backgrounds. The motivation to perform multiple comparisons has only increased as a result of progress in the areas of personalized therapies and discovery of biomarkers (Shyr & Liu, 2013). Some of the most popular tools for AS analysis, like rMATS (Shen et al., 2014), MAJIQ (Green et al., 2018), and MISO (Katz et al., 2010), offer a combined solution for quantification and comparison across pre-aligned BAM files generated with other tools such as STAR aligner (Dobin et al., 2013) (Supplemental Figure 2B). While these are all highly accurate tools for AS analysis, they carry the burden of unnecessary repeated quantifications, when only the comparisons should be repeated. To expedite data analysis, SpliceCore offers an alternative software design, by pulling one-time alignment and quantification with SpliceTrap, allowing for fast and scalable statistical modeling of AS comparison using SpliceDuo. Benchmarking to open-source competitors demonstrated that SpliceCore performs in a significantly larger search space (Supplemental Figure 2C) with outstanding speed (Supplemental Figure 2D-G), scalability (Supplemental Figure 2H-J) and accuracy (Supplemental fig. 3K-O), thereby accelerating value extraction from RNA-seq data.

### SpliceLearn is an AI/ML method for predicting functional SSO binding sites

Previous methods for antisense oligonucleotide binding predictions were designed to predict RNA down-regulation instead of splicing changes (Giddings et al., 2002) or focused on SF binding perturbations without informing whether such binding affects the AS outcome (Bjørnholt Grønning et al., 2020). In recent years, novel methods to predict the effect of single nucleotide variants (SNV) on the AS outcome have been developed (Cheng, Yen, et al., 2019; H. Y. Xiong et al., 2014). Like SSOs, SNVs can also perturb RNA sequences to change AS. Current methods to predict the effect of SNVs on splicing regulation can be used to identify SSO binding sites in RNA sequences (H. Y. Xiong et al., 2014). However, such methods do not inform the identity of specific SFs potentially blocked by binding perturbations. Informing the identity of prospective SFs blocked by SSO can help accelerate drug development by increasing the biological interpretability, assisting experimental design, and enabling integration with other data types, such as CLIP-seq or genomics, to better understand SSO mechanism and SF involvement in disease progression. SpliceLearn innovates by offering the combined benefit of robust AS outcome predictions with high biological interpretability using machine learning.

### SpliceLearn was trained and validated using massively parallel splicing reporter assays

To train and test SpliceLearn, we utilized data from a massively parallel splicing minigene reporter assay (MFASS) that quantifies the effect of SNVs on AS outcomes, therefore mimicking the impact of SF-blocking by SSOs (Cheung et al., 2019). To develop predictive models with high interpretability, we estimated the differential effect of SNVs in the binding motif scores of 141 SFs derived from three different methods (Paz et al., 2014; Ray et al., 2009; X. Wang et al., 2011). We combined SFs into 83 non-exclusive clusters (SFCs), based on spliceosomal functional annotations (Table S2) and SF-SF binding probabilities (Akerman et al., 2015). Each SFC was composed of physically interacting SFs attributed to a given splicing-related function such as membership to spliceosomal subcomplexes (e.g. U1 snRNP), RNA-binding specificity (e.g. AG binding), regulatory outcome (e.g. repressor) and others (Table S2). Since SFs perform multiple functions in splicing regulation, they were non-exclusive and permitted to appear in more than one SFC. The advantage of using SFCs as predictive features in lieu of SFs scores is that they reduced matrix sparsity and zero-inflation, while capturing functional, regulatory, and evolutionary aspects of the spliceosome. By grouping SFs into SFCs, we provided a more intuitive context for biological interpretation.

We tested the ability of each independent SFC to differentiate between “positives” and “negatives” using the Wilcoxon test with Holm-Sidak p-value adjustment (W. Guo & Romano, 2007). We performed the analysis in exons, upstream and downstream introns independently. We observed that the three sequence types were characterized by different subsets of significant SFCs (Table S2). For instance, the “distance to splice site” SFCs were highly significant in introns (*adj*.*p-val<5*.*06E-11*) but not exons (*adj*.*p-val*≤0.356). This observation was expected and agrees with many studies showing intronic sequences around the splice sites to be enriched with SF binding sites important for AS regulation (van Nostrand et al., 2016; Z. Wang & Burge, 2008; Yee et al., 2019; Yeo et al., 2007). In addition, SR proteins and activators were highly significant SFCs in exons (*adj*.*p-val*≤2.14E-08 and *adj*.*p-val*≤4.91E-05, respectively) but not introns (0.003≤*adj*.*p-val*≤0.98), consistent with several studies describing the role of SR proteins as splicing activators that bind exonic splicing enhancers (Jeong, 2017; Lin & Fu, 2007; J. Wang et al., 2005). In upstream introns, SF binding sites such as U-rich (*adj*.*p-val*≤6.64E-07) and CG-rich (*adj*.*p-val*≤2.1E-06) motifs were highly significant. Interestingly, CG-rich introns have been associated with weak 3’ splice sites and polypyrimidine tracts, important for the regulation of alternative exons (Murray et al., 2008; E. J. Wagner & Garcia-Blanco, 2001). These polypyrimidine tracts, which are intrinsically uridine-rich, are known for attracting several SFs (Barreau et al., 2006), including members of the A complex such as U2AF2, known for its crucial role in 3’ splice site recognition and exon inclusion (Graveley et al., 2001; R. Singh et al., 2000; Warnasooriya et al., 2020). Accordingly, the “A complex” SFC was more significant in upstream (*adj*.*p-val*≤0.0006) and downstream introns (*adj*.*p-val*≤3.62E-13) than exons (*adj*.*p-val*≤0.756). Other highly significant SFCs in downstream introns were “repressors” (*adj*.*p-val≈0*) and members of the hnRNP family (*adj*.*p-val≈0*) known to interact with intronic splicing silencers to inhibit exon inclusion (Geuens et al., 2016). In addition, SFCs corresponding to hnRNP binding motifs were highly significant, including UG-rich (*adj*.*p-val≈0)*, CU-rich (a*dj*.*p-val*≤1.80E-14) and CA-rich (a*dj*.*p-val*≤4.11E-13) elements. The U1 snRNP SFC was also highly significant in the downstream intron (*adj*.*p-val*≤1.84E-10). In summary, a total of 2, 5 and 14 SFCs showed *adj*.*p-val*≤1.0E-06 in exons, upstream and downstream introns, respectively. This distribution suggested that the base information for AI/ML model development was particularly rich in the downstream introns.

We then trained SpliceLearn using the MFASS data for exonic, upstream, and downstream intronic sequences. We tested six different types of AI/ML classifiers and in all of the cases, the best performing model type were XGboost trees, an implementation of gradient boosted decision trees designed for speed and performance (Sheridan et al., 2016). Prominently, SpliceLearn-down was the top performing model with an AUC of 0.95, followed by SpliceLearn-up with an AUC of 0.88 and finally SpliceLearn-exon with AUC of 0.60. The sensitivity and specificity were high for both SpliceLearn-downstream (0.92 and 0.93 respectively) and SpliceLearn-upstream (0.84 and 0.75) although for SpliceLearn-exon only the specificity was relatively high (0.71) while the sensitivity was poor (0.40) (Supplemental Figure 3). Notably, both intronic SpliceLearn classifiers clearly outperformed the exonic one, suggesting that it would be easier to predict productive SSO binding sites in introns vs exons. This could be due the fact that, unlike exons, introns are not subjected to protein-coding constraints, thus regulatory information may be easier to identify in introns using AI/ML (Z. Wang & Burge, 2008). In addition, intronic regions near the splice sites are preferable for SSO targeting compared to exons, since introns are only present in nuclear pre-mRNA, while exons are both present in pre- and mRNA, potentially increasing the chances of off-target effects. For example, the SSO Nusinersen that treats Spinal Muscular Atrophy targets the intron downstream of exon 7 in the *SMN2* pre-mRNA, to block binding of the splicing repressor hnRNPA1 (R. N. Singh & Singh, 2018). To test SpliceLearn with an independent data source, we used the Vex-seq dataset, which consists of 1226 qualifying variants that were experimentally identified to have impact on pre-mRNA splicing using a high-throughput reporter assay (Adamson et al., 2018). As a result, The SpliceLearn-down model classified VexSeq data with an AUC of 0.98, SpliceLearn-up with and AUC of 0.86 and SpliceLean-e with AUC of 0.66 (Supplemental Figure 3). These results confirm the usefulness of both upstream and downstream intronic SpliceLearn models to predict SF-binding perturbation positions critical for AS regulation and useful for SSO targeting.

### SpliceLearn balances performance and interpretability

We compared SpliceLearn’s predictive accuracy to that of two competitive methods, SPANR, a tool to predict SNVs effect on AS outcome, which has been used before for SSO design,(H. Y. Xiong et al., 2014) and a more recent method called MMSplice (Cheng, Yen, et al., 2019), which was previously shown to outperform other equivalent tools by winning the CAGI 5 splicing challenge (Cheng, Çelik, et al., 2019). The overall performance of the three methods was comparable, with MMSplice showing slightly better AUC for upstream introns (MMsplice=0.88, SpliceLearn=0.86, SPANR=0.77) and SpliceLearn matching MMsplice’s AUC for the downstream intron (SpliceLearn=0.95, MMSplice=0.95, SPANR=0.89) (Supplemental Figure 4).

While SpliceLearn performance was comparable to competitive tools, it showed significant advantage in interpretability with more balanced sensitivity/specificity overall. SpliceLearn was developed using tree-based learning (i.e. XGboost trees), an AI/ML methodology that is inherently more interpretable because it enables the investigation of predictive features content and relationships. In addition, tree-based methods output probabilistic quantities (in a scale from 0 to 1) which are intuitive to users (Azodi et al., 2020). We computed the sensitivity/specificity trade-off of SpliceLearn, MMsplice and SPANR using the Youden index, a summary measure that enables the selection of an optimal threshold values for predictive model (Fluss et al., 2005). SpliceLearn’s optimal threshold was 0.5 for both -up and -down intronic models, with Youden indexes of 0.55 and 0.84, and sensitivity/specificity tradeoffs of 0.82/0.73 and 0.91/0.93 respectively (Supplemental Figure 4C). These results indicate that SpliceLearn performs with balanced sensitivity/specificity and an intuitive threshold that appropriately represents the midpoint of the tree-based probability distribution. In contrast, MMsplice and SPANR scored with relatively lower Youden indexes (upstream introns 0.4 and 0.27 downstream introns 0.36 and 0.49) leading to less balanced sensitivity/specificity. For instance, the sensitivity/specificity tradeoff at the Youden inflexion point were 0.92/0.48 and 0.79/0.48 for MMsplice and SPANR, respectively, in the upstream intron, and 0.96/0.53 and 0.89/0.47, respectively, in the downstream intron. In addition to a strong bias towards sensitivity, the selected optimal thresholds for these tools (−0.1 to 0, respectively) seemed subjective and not representative of the training data distribution or positive/negative ratios (Supplemental Figure 4B).

### Identification of most predictive SFs

To deliver SSO target discovery using SpliceLearn, it is necessary to backtrack the identity of SFs to be displaced by SSOs. Knowledge of these specific SSOs is not only crucial for the biological interpretability, but also to facilitate experimental drug design and integration with other data types (e.g. CLIP-seq). To this aim, we derived a feature selection approach based on Shapley additive explanation analysis (SHAP). SHAP is a game theory approach to estimate the importance of specific predictive features to the overall performance of AI/ML models (Lundberg et al., 2019).

Tree learning algorithms like XGboost build several decision trees by bootstrapping the training data. For a given predictive feature, the SHAP value is the average marginal contribution of this feature value across all possible decision trees. A key advantage of SHAP over other feature importance inference methods, is that it is unaffected by the order in which features are randomly chosen by tree-models, thus it is a robust tool for the interpretation of the primary information driving predictive efficiency in AI/ML (Lundberg et al., 2019). To complement the SHAP analysis we also performed an additional feature prioritization analysis where the data considered at each bootstrap sample (called bag data) are considered to learn a classifier and the remaining training data are considered as out-of-bag data (OOB) (Breiman, 2001).

We applied OOB and SHAP to weigh the contribution of every SFC to the SpliceLearn-upstream and SpliceLearn-downstream models. OOB/SHAP scatter plot revealed the “distance to splice site” information as a major driver of predictability, whereby positions closer to the splice sites show greater potential of AS alterations (Supplemental Figure 5A, C). Strikingly, this observation suggests that the position of perturbing factors (i.e. SNVs, SSOs) relative to the splice sites is sufficient to explain much of the AS outcome. Despite its strong predictive power, “distance to splice sites” as predictive feature does not provide interpretability regarding the role of specific SFs, because distance is consistent at a single nucleotide for every SF, rather than subsets, as in the case of all other SFCs. To avoid the dominating effect of “distance to splice site” and allow SHAP to best prioritize interpretable SFCs, we retrained SpliceLearn-upstream and -downstream, only this time after without any “distance to splice site” information (Supplemental Figure 5B, D). As a result, we observed that despite a proportional reduction in predictive efficacy, SpliceLearn-downstream still performed well with an AUC of 0.80, while SpliceLearn-upstream showed more borderline, yet clear predictive power with an AUC of 0.67 (Supplemental Figure 3C).

### Validation of most predictive SFs

SHAP/OOB values can be estimated for every predictive feature at every tested data point (i.e., nucleotide), and therefore we can estimate SHAP/OOB values per SFC at each nucleotide position to identify the most important SFs driving predictive accuracy of every putative SSO binding site. Furthermore, each SF within an SFC has percentile scores from each of the three methodologies used to measure SF binding (Paz et al., 2014; Ray et al., 2009; X. Wang et al., 2011), which allows us to rank SFs within the SFC. We used these datapoints to find the most predictive SFs within each SFC. While these scores were calculated from computational and *in-vitro* approaches, *in-vivo* approaches such as eCLIP can be used to confirm if an SF occupies a specific location in the genome (in the cell type of interest). We used eCLIP data from ENCODE (Encode Project Consortium, 2012) to evaluate the correspondence between measured predictivity in SpliceLearn and evidence of binding *in-vivo*. We compiled SpliceLearn predictions from 2,124 introns and started by determining the most predictive SFs in the SFCs in the top 25% (based on ranked SHAP values) with the best binding scores (percentile score > 50) at each position where the splicing effect probability was at least 0.5 (as previously established in Supplemental Figure 4C). We found that the sites where several SFs are predictive to influence splicing outcomes are highly enriched for eCLIP peaks at those same sites (Supplemental Figure 5E-F). Using the odds ratio as a metric, we then proceeded to test a wide range of cutoffs for each variable to determine the set of conditions that best enriches the SpliceLearn hits for eCLIP peaks for each individual SF. Generally, we found that using a highly stringent cutoff for the splicing effect probability allowed for the most robust enrichment with the eCLIP data (Supplemental Figure 7G). While this stringent cutoff for splicing effect probability enriches the hits for eCLIP peaks, it is likely that this will be too stringent when looking to identify sites to test with SSOs. Overall, we have shown that SpliceLearn provides much better biological interpretability than alternative methods, which is crucial for the development of novel therapeutics. Specifically, we have seen that the predictions made by SpliceLearn can be traced back to one or multiple SFs, and that these predicted SF binding sites are enriched for eCLIP peaks in independent datasets, confirming the binding of those SFs at those locations.

### Western Blot

Total cell lysates were prepared from breast cancer cell lines using RIPA or NP-40 buffer in the presence of protease inhibitor. Nuclear, cytoplasmic and membrane protein extracts were obtained according to the Subcellular Protein Fractionation Kit for Culture Cells (Thermo Scientific #78840) manufacturer instructions. About 10ug of total protein was separated in 4-15% gradient gel. The membrane bound specific protein bands were detected using specific primary antibodies (phSMAD2/3 (Cell Signaling) 1:1000, SMAD2/3 (Cell Signaling) 1:1000, TGFβ Receptor I (EMD Millipore), 1:500, Tubulin (Cell Signaling) 1:1000, Actin (Sigma Aldrich) 1:5000 and H3 (Cell Signaling) 1:1000 followed by incubation with HRP conjugated secondary antibody and chemiluminescence detection (Biorad) according to the manufacturer’s instruction. Tubulin was used as a loading control.

## Notes

### Competing Interest Statement

Alyssa D Casill, Miguel A Manzanares, Paulina Zheng, Adam Geier, Kendall Anderson, Vanessa Frederick, Shaleigh Smith, Sakshi Gera, Robin Munch, Mahati Are, Priyanka Dhingra, Gayatri Arun, and Martin Akerman were either previously or are currently employed by Envisagenics.

